# Identification of a minimal *Alu* domain required for retrotransposition

**DOI:** 10.1101/2024.12.16.628748

**Authors:** John B. Moldovan, John Yin, John V. Moran

## Abstract

*Alu* elements are primate-specific retrotransposon sequences that comprise ∼11% of human genomic DNA. *Alu* sequences contain an internal RNA polymerase III promoter and the resultant *Alu* RNA transcripts mobilize by a replicative process termed retrotransposition. *Alu* retrotransposition requires the Long INterspersed Element-1 (LINE-1) open reading frame 2-encoded protein (ORF2p). Current models propose that *Alu* RNA binds to signal recognition particle proteins 9 and 14 (SRP9/14) and localizes to ribosomes, which allows *Alu* to ‘hijack’ L1 ORF2p. Here, we used HeLa cell-based retrotransposition assays to define a minimal *Alu* domain necessary for retrotransposition. We demonstrate that *Alu* transcripts expressed from a cytomegalovirus (CMV) RNA polymerase II promoter can efficiently undergo retrotransposition. The use of an external CMV promoter to express *Alu* RNA allowed us to construct separation-of-function mutations to examine the effects of large deletions within the *Alu* sequence on retrotransposition. Deletion mutagenesis demonstrated that a 46 nucleotide (nt) domain located at the 5′ end of the *Alu* RNA transcript is necessary for *Alu* retrotransposition. Consistent with current models, the 46 nt 5′ *Alu* domain associates with SRP9/14 in HeLa-HA cell extracts and can promote a single round of retrotransposition in HeLa-HA cells. We propose that the 46 nt 5′ *Alu* domain forms a discrete structure that allows for SRP 9/14 binding and ribosomal association, thereby allowing the *Alu* poly(A) tract to compete with the L1 poly(A) tail for ORF2p RNA binding to mediate its retrotransposition.

## INTRODUCTION

*Alu* elements are a class of Short INterspersed Elements (SINEs) present at high copy numbers in all primate genomes (1). *Alu* elements are related to the *7SL* RNA gene and encode a short non-coding RNA polymerase (pol) III transcript (2). *Alu* elements mobilize throughout the genome by a replicative process termed retrotransposition (3), which requires the Long INterspersed Element-1 (LINE-1 or L1) retrotransposon open reading frame 2 encoded protein (ORF2p) (4). *Alu* is the most active retrotransposon in the human genome, and it is estimated that a new germline *Alu* insertion occurs in ∼1 out of 40 human births (5,6). Over the course of primate evolution, *Alu* retrotransposition has resulted in the accumulation of ∼1.1 million Alu elements in the human genome (7–9).

The *7SL* RNA is a major structural component of the signal recognition particle (10,11) and has given rise to several primate, rodent, tree shrew, and hagfish SINE families (12,13). The *Alu* SINE family is specific to primates, appearing within the last 65 million years (14). Alu elements are classified into subfamilies based on diagnostic sequence identifiers, pairwise sequence divergence, and evolutionary age (1). The *Alu*Y subfamily is the youngest *Alu* subfamily and is still active in the human genome.

An active human *Alu*Y element is approximately 280 nucleotides long and consists of two highly similar *7SL Alu* domain monomers linked together by a short adenosine (A)-rich sequence (**Fig 1A**) (15). The *Alu* 5′ left *Alu* domain monomer contains a pol III A and B box promoter that directs the initiation of *Alu* transcription at or near the first *Alu* nucleotide (16). The 3′ end of the *Alu* sequence is characterized by a poly(A) tract; poly(A) tract lengths of 40 or more nucleotides are associated with higher rates of retrotransposition activity (17). Younger human *Alu*Y subfamily members typically contain longer 3′ poly(A) sequences than older *Alu*S and *Alu*J subfamilies (18).

**Figure 1:**
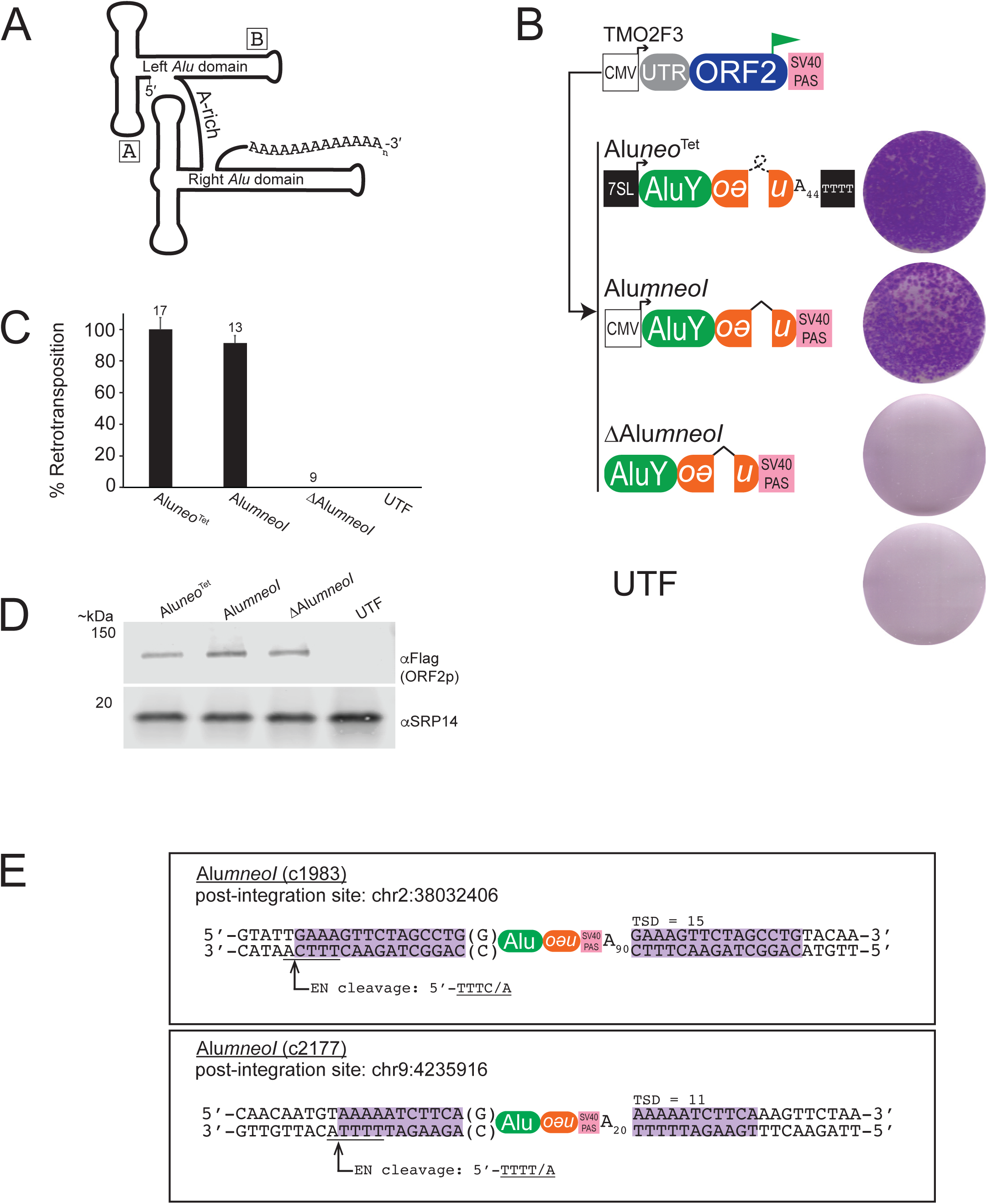
Retrotransposition of an *Alu*Y element transcribed by an RNA pol II CMV promoter. *(A) Schematic of a consensus human AluY element*. *Alu*Y is ∼282 nt in length and consists of two *7SL* RNA *Alu* domain monomers (*left* and *right*) connected by an A-rich linker and ends in a variable length 3′ encoded poly(A) tract. The approximate locations of the pol III A and B boxes are indicated. *(B) Results of AluY retrotransposition assays*. HeLa-HA cells were co-transfected with TMO2F3 and the indicated *Alu*Y plasmid. TMO2F3 expresses ORF2p with a carboxyl terminus 3XFLAG tag (green flag). ORF2p expression is augmented by a CMV promoter (white square), the native L1 5′ UTR (grey oval), and an SV40 PAS (pink square). The *Aluneo*^Tet^ retrotransposition marker (orange oval) contains a backwards copy of the neomycin phosphotransferase gene interrupted by a self-splicing group I intron (loop) that is in the same transcriptional orientation as the *Alu* sequence. A 44 bp encoded poly(A) tract follows the *neo*^Tet^ sequence. *Aluneo*^Tet^ expression is augmented by a *7SL* gene enhancer (black square) and a sequence of four consecutive thymidine residues (black square) located downstream of the 44 bp poly(A) tract. The *AlumneoI* retrotransposition marker (orange oval) contains a backward copy of the neomycin phosphotransferase gene interrupted by a γ-globin intron (^) that is in the same transcriptional orientation as the *Alu* sequence. *AlumneoI* expression is augmented by a CMV promoter (white square) and an SV40 PAS (pink square) located after the *mneoI* indicator. Displayed next to each plasmid construct is a single well of a representative six-well tissue culture plate from retrotransposition assays. UTF = untransfected. *(C) Quantification of AluY retrotransposition assays.* The X-axis indicates *Alu* expressing construct co-transfected with TMO2F3 and the Y-axis indicates average percent (%) retrotransposition normalized to *Alu*neoTet from (n) independent biological replicates for each transfection condition. Error bars indicate standard deviation; (n) number of biological replicates indicated above the error bars. *(D) ORF2p-Flag expression levels.* Western blots of HeLa-HA lysates co-transfected with TMO2F3 and the *Alu* retrotransposition construct (top of lane). The antibody used for detection is indicated on right. MW markers are indicated on the left. Western blot experiments were repeated twice with similar results. *(E) Structure of de novo CMVAlumneoI insertions (c1983, top) and (c2177, bottom).* Genomic insertion site locations are based on human reference genome sequence T2T CHM13v2.0/hs1. The *AlumneoI* insertions contain typical TPRT structural hallmarks indicative of L1 ORF2p-mediated retrotransposition (see main text). Target site duplications are highlighted in magenta. The ORF2p EN target site is underlined; the arrow and “/” indicates the L1 EN cleavage site. Parentheses indicate untemplated nucleotides.

According to current models (4,19), efficient *Alu* transcription requires intact A and B box promoter elements, as well as genomic DNA sequences at the 5′ flank of the *Alu* element that enhance transcription (16,20,21). *Alu* transcription terminates at a sequence of four or more thymidine nucleotides in the 3′ flanking genomic DNA downstream of the *Alu* 3′ poly(A) tract. After transcription, *Alu* RNA binds to the signal recognition particle proteins 9 and 14 (SRP9/14) to form a ribonucleoprotein particle (RNP) (22–24). The *Alu* RNP then is hypothesized to localize to ribosomes, which allows the *Alu* RNA poly(A) tail to compete with the L1 RNA poly(A) tail to bind ORF2p (4,19,25,26). Upon ORF2p binding, *Alu* RNA presumably departs the ribosome and enters the nucleus, where the *Alu* RNA is copied and inserted into a new genomic location by the ORF2p-mediated process of target-site primed reverse transcription (TPRT) (4,27).

*Alu* RNA sequences required for retrotransposition include the internal A and B box pol III promoter (16) and a 3′ poly(A) tract, which is necessary to bind L1 ORF2p (4,26). Here, we sought to define a minimal *Alu* RNA sequence required for retrotransposition. Evolutionary comparisons of *Alu* and other *7SL*-derived SINEs from mammals and hagfish indicate that the *Alu* domain structure is highly conserved among all *7SL*-derived SINEs (13,28). In addition, the *Alu* domain RNA structure is crucial for SRP9/14 binding and ribosome interaction; mutations that decrease *Alu* RNA affinity for SRP9/14 binding negatively impact *Alu* retrotransposition (25,29).

Here, we used a well-established *Alu* retrotransposition assay (4), in conjunction with deletion mutagenesis, to define a minimal *Alu* domain necessary for retrotransposition. We first engineered vectors that faithfully express *Alu* RNA using an external pol II cytomegalovirus (CMV) immediate early promoter and demonstrate that these *Alu* RNAs can efficiently undergo a single round of retrotransposition in Hela-HA cells. The ability to bypass the requirement for the A and B box promoter elements then allowed us to create separation-of-function deletion mutations to identify a minimal Alu domain required for retrotransposition. Our results demonstrate that a discrete 5′ *Alu* domain RNA consisting of 46 nucleotides (nts) can associate with SRP9/14 in HeLa-HA cell extracts and efficiently retrotranspose in HeLa-HA cells. Thus, we have identified a mini-*Alu* domain required for retrotransposition in HeLa-HA cells.

## MATERIALS & METHODS

### Cell Culture Conditions

The HeLa-HA strain was a kind gift from the laboratory of Dr. Astrid Roy-Engel (New Orleans, LA, USA) and was extensively characterized in a previous study (30); it was the only cell line used in this study. HeLa-HA cells were grown in high-glucose DMEM (Gibco) supplemented with 10% Fetal Bovine Serum (FBS) (Gibco), 100 U/mL penicillin-streptomycin (Invitrogen), and 0.29 mg/mL L-glutamine (Gibco) as described previously (30). All cells were maintained at 37°C with 7% CO_2_. We regularly screened cells for the absence of *Mycoplasma spp*.

### Plasmids

Oligonucleotide sequences used in this study and cloning strategies are available upon request. All human L1 plasmids contain the indicated fragments of L1.3 DNA (a retrotransposition-competent human L1, GenBank accession no. L19088 (31)) cloned into the pCEP4 (Invitrogen) episomal vector unless indicated otherwise. Plasmid DNAs were prepared using a Midiprep Plasmid DNA Kit (Qiagen).

<u>TMO2F3</u>: is a pCEP4-based plasmid that contains the L1.3 native 5′ UTR and ORF2 sequences; the 3’ end of ORF2 contains a sequence that encodes a 3XFLAG epitope-tag (32). L1 ORF2 expression is augmented by the pCEP4 CMV promoter and SV40 polyadenylation signal sequence (PAS).

TMO2F3(D702A): is derivative of TMO2F3 that contains a D702A (GAC to GCU) missense mutation in the RT active site of L1.3 ORF2 (32).

*Aluneo*^Tet^: expresses an *Alu*Y element that is marked with the *neo*^Tet^ retrotransposition reporter cassette. The reporter was subcloned upstream of a 44 nucleotide (nt) long poly(A) tract (4,33). *Alu* expression is augmented by the *7SL RNA* enhancer located upstream of the *Alu* sequence and an RNA polymerase III terminator located after the 44nt poly(A) sequence. This construct was provided by Dr. Thierry Heidmann (Villejuif, France).

*<u>AlumneoI</u>*: contains the *Alu*Y sequence from *Aluneo*^Tet^, which has been cloned into the pCEP4 vector (Invitrogen) downstream of the CMV promoter transcription start site. The *Alu*Y is marked with the *mneoI* retrotransposition reporter cassette (34). *Alu*Y expression is augmented by the pCEP4 CMV promoter and an SV40 PAS located after the *mneoI* indicator.

Δ *AlumneoI*: is a derivative of *AlumneoI* that contains a deletion of the entire CMV promoter sequence.

<u>EGFP*mneoI*</u>: is a pCEP4 based vector containing the enhanced green fluorescent protein *(EGFP) ORF* sequence from plasmid pEGFP-C3 (Clontech) followed by the *mneoI* retrotransposition indicator cassette. The *EGFP* and *mneoI* sequences are cloned in between the pCEP4 CMV promoter and SV40 polyadenylation sequences (30).

EGFP*AlumneoI*: is a derivative of EGFP*mneoI* that contains the *Alu*Y sequence from *Alu*neo^Tet^ inserted between the *EGFP* ORF and the *mneoI* indicator cassette of EGFP*mneoI*.

SA86*mneoI*: is a derivative of *AlumneoI* where the *Alu*Y sequence has been replaced by SA86.

SA46*mneoI*: is a derivative of *AlumneoI* where the *Alu*Y sequence has been replaced by SA46.

SA39*mneoI*: is a derivative of *AlumneoI* where the *Alu*Y sequence has been replaced by SA39.

SA20*mneoI*: is a derivative of *AlumneoI* where the *Alu*Y sequence has been replaced by SA20.

*Alu*Y46*mneoI*: is a derivative of *AlumneoI* where the *Alu*Y sequence has been replaced by *Alu*Y46.

CMV*mneoI*: is a derivative of *AlumneoI* that contains a deletion of the entire *Alu*Y sequence.

<u>EGFP*Alumneoi*ColE1</u>: is a derivative of EGFP*AlumneoI* that contains a ColE1 bacterial origin of replication located between the *mneoI* indicator and the SV40 PAS. EGFP*AlumneoIK7* was constructed by inserting the *Ngo*MIV-*Bam*H1 fragment spanning the *mneoI*-ColE1 sequence from pJM140/L1.3/delta2K7 (35,36) into EGFP*AlumneoI*.

CMV*mneoI*ColE1: is a derivative of CMVmneoI that contains a ColE1 bacterial origin of replication between the *mneoI* indicator and the SV40 PAS and was constructed by inserting the *Ngo*MIV-*Eco*RV fragment from EGFP*AluneoK7* into CMV*mneoI*.

*Alublast*: is a derivative of *AlumneoI* that contains a blasticidin retrotransposition indicator (37) in place of the *mneoI* indicator.

Δ*Alublast*: is a derivative of *Alublast* that contains a deletion of the entire CMV promoter sequence.

### *Alu* Retrotransposition Assays

For all *Alu* RNA retrotransposition assays (4,38), approximately 2-3×10^5^ HeLa-HA cells were plated per well in 6-well tissue culture plates (Corning). The next day, the HeLa-HA cells were transfected with 0.5 μg of “driver” plasmid (TMO2F3) + 0.5 μg of “reporter” plasmid (*Aluneo*^Tet^ or other *Alu* reporter plasmids) using 3 μL FuGENE® HD (Promega). The following day (∼24 hours), the transfection media was removed, and fresh media was added to each well. Three days post-transfection, cells were grown in the presence of G418 (400 μg/mL) to select for *Alu* retrotransposition events; the media was replaced with fresh G418-containing media approximately every-other day throughout the selection period. Transfection efficiency was assessed ∼48-72 hours after transfection by monitoring ORF2p-FLAG expression using Western blot analysis. After approximately 12 days of selection, cells were washed with 1x phosphate buffered saline (PBS), fixed to plates using (2% formaldehyde, 0.2% gluteraldehyde, in 1× PBS, pH 7.4), and then stained with crystal violet to visualize G418-resistant colonies. To quantify retrotransposition, G418-resistant colonies were counted visually, and the average number of G418-resistant foci were determined for each transfection condition, which was extrapolated from the foci count from a single well from a 6-well tissue culture plate from three or more independent experiments.

### Characterization of Insertions by Inverse PCR

Inverse PCR was performed as described previously (37,39,40) with minor modifications. Briefly, ∼2×10^4^ HeLa-HA cells were plated into single wells of a six well plate. After ∼12 hours, HeLa-HA cells were transfected using 3 μL of FuGENE 6 (Promega) and 0.5 μg *Alu* reporter plasmid + 0.5 μg TMO2F3 plasmid DNA per well of a six well plate as noted above. Three days post-transfection, the cells were subjected to G418 selection (400 μg/mL). After 12 days, individual G418-resistant foci were isolated and expanded in 6-well plates until wells were ∼80% confluent. Five micrograms of genomic DNA extracted from the expanded G418-resistant foci (Qiagen) were digested to completion with *Ssp*I-HF (NEB) in a 100 μL reaction volume. The restricted DNA was concentrated by ethanol precipitation and then ligated using 8 μL (3200U) of T4 DNA ligase (NEB) in a volume of 600 μL at 16°C overnight. The ligated DNA was precipitated using a 0.6x volume of isopropanol, washed with 70% ethanol, and resuspended in 50 μL of water. Five microliters of the ligated DNA (∼100-200 ng) were used in the primary PCR reaction (Q5 Hot Start High-Fidelity 2X Master Mix, NEB) in a 25 μL reaction volume containing 500 nM of oligonucleotide primers (NEO210AS: 5′-GACCGCTTCCTCGTGCTTTACG-3′) and (NEO742S: 5′-TCATTCAGGGCACCGGACAG-3′). Samples were amplified using the following thermocycler conditions: one cycle of 98°C for 30 sec, then 30 cycles of 98°C for 10 sec, 68°C for 30 sec, and 72°C for 4 min, followed by a final extension step at 72°C for 5 min. The resultant PCR products (5 μL) were used in a second PCR reaction using the same conditions with the nested oligonucleotide primers (NEO148AS: 5′-CGAGTTCTTCTGAGGGGATTTG-3′) and (NEO1808S: 5′-GCGTGCAATCCATCTTGTTCAATG-3′). PCR products from the second amplification were purified using a gel extraction kit (NEB T1020). Purified DNA from excised gel slices was cloned into a pCR4Blunt-TOPO vector using a Zero Blunt TOPO PCR Cloning Kit for Sequencing (Thermo Fisher Scientific, 450159) and subjected to Sanger DNA sequencing using M13 forward and M13 reverse primers (Eurofins Genomics). Sequences flanking the *Alu* insertion were used as probes in BLAT search at the UCSC genome browser (http://genome.ucsc.edu) T2T CHM13v2.0/hs1 (41) to determine the insertion site location (42).

### Characterization of insertions using recovery method

The recovery of L1 integration events as autonomously replicating plasmids in *E. coli* was conducted as described previously (35,36,43,44) with minor modifications. Briefly, ∼2×10^5^ HeLa-HA cells were plated into single wells of a six well plate. After ∼12 hours, HeLa-HA cells were co-transfected using 3 μL of FuGENE HD (Promega) with 0.5 μg of recovery vector (*i.e.*, EGFP*AlumneoI*ColE1, CMV *mneoI*ColE1) and 0.5 μg TMO2F3 plasmid DNA per well of a six well plate. Three days post-transfection, the cells were subjected to G418 selection (400 μg/mL). After 12 days, the G418-resistant foci were pooled, and genomic DNA was extracted using a genomic DNA extraction kit (Qiagen). Five micrograms of genomic DNA then were digested overnight at 37°C with 5 μL of *Ssp*I (NEB), or *Sca*I (NEB) in a 100 μL reaction volume. Digests then were heat-inactivated, the restricted DNA was ligated by adding 8 μL (3200U) of T4 DNA ligase (NEB) directly to the digested DNA in a volume of 600 μL. The ligation reactions were allowed to proceed at 16°C overnight. The ligation reactions then were heat inactivated at 60 °C for 20 minutes and the resultant ligated DNA was concentrated using an Amicon YM-100 filter column to a final volume of 50-60 µL. Approximately 20 µL of the concentrated DNA (∼800 ng-1000 ng) was used to transform 100 µL of XL-10 Gold ultra competent cells (Agilent) (genotype: Tet^r^Δ(*mcrA*)*183* Δ(*mcrCB-hsdSMR-mrr*)*173 endA1 supE44 thi-1 recA1 gyrA96 relA1 lac* Hte [F *proAB lacI*^q^*Z*Δ*M15* Tn*10* (Tet^r^) Amy Cam^r^]), which then were incubated overnight at room temperature with shaking at 225 rpm. The transformed XL-10 Gold cells were then plated on 15 cm (corning) kanamycin (25 µg/mL) agar plates and incubated at 37 °C for 18-14 hours. Plasmid DNA was extracted from individual kanamycin-resistant clones and then analyzed by whole plasmid sequencing (Eurofins) to determine the insertion sequence and flanking genomic DNA regions. The sequences flanking the insertions were used as probes in BLAT search (41) at the UCSC genome browser (http://genome.ucsc.edu) T2T CHM13v2.0/hs1 (42) to determine the insertion site location.

### *Alu* RNA pulldown assays

HeLa-HA cells were cultured in 15 cm dishes (Corning) and grown to confluency. The cells then were washed with 1x PBS, scraped into 15 mL centrifuge tubes, centrifuged at 1000×g for 5 minutes, and the resultant cell pellets were frozen at –80°C. Frozen cell pellets were thawed and lysed (1 mL of lysis buffer was used per ∼100 mg HeLa-HA cell pellet) in lysis buffer (150 mM KCl, 1 mM MgCl_2_, 25 mM Tris-HCl [pH 7.5], 1 mM DTT, 0.5% NP-40, and 1× protease inhibitor EDTA-free mixture [Roche]) on ice for 30 minutes. Whole cell lysates then were centrifuged at 12,000×g for 15 minutes at 4°C. The supernatant (250 µL) was then transferred to a fresh 1.5 mL tube and incubated with 50 µL of washed streptavidin T1 Dynabeads (Invitrogen) for 30 minutes at room temperature to pre-clear the lysates. The pre-cleared lysates were then transferred to a fresh 1.5 mL and stored on ice.

The 5′-biotin labeled *Alu* RNA oligonucleotides (Table S1) were obtained from Integrated DNA Technologies (IDT). To attach *Alu* RNA oligonucleotides to streptavidin T1 Dynabeads, 20 µL of stock RNA oligo (100 µM) was diluted using 250 µL of binding buffer (5 mM Tris-HCL [pH7.5], 0.25 mM EDTA, 0.5M NaCl) to a final concentration of ∼8 µM, and then was combined with 50 µL (0.5 mg) of washed streptavidin T1 Dynabeads in a 1.5 mL microcentrifuge tube. The RNA-bead mixture was incubated for 15 minutes at room temperature with nutation to bind RNA to the streptavidin beads. The beads were then washed 3 times with binding buffer.

For RNA pulldowns, 250 µL of pre-cleared HeLa-HA cell lysate was added to the washed RNA-streptavidin bead complexes and incubated at room temperature for 60 minutes with nutation. The beads were then washed 3 times with 500 µL of cold wash buffer (50 mM Tris-HCL [pH7.5], 150 mM NaCl, 0.05% NP-40, 1 mM DTT, and 1× protease inhibitor EDTA-free mixture [Roche]). RNA-protein complexes were eluted from the washed beads by incubating with 60 µL of 2× Laemmli buffer at 95 °C for 5 minutes. Approximately 5 – 10 µL of the eluted fraction was then analyzed by SDS-PAGE.

### Western Blotting and Primary Antibodies

Standard western blotting procedures were used for protein analysis (45). Blots were analyzed using an Odyssey CLx (LI-COR) using the Image Studio software (version 3.1.4, LI-COR). The αSRP9 (11195; 1:1,000 dilution) and αSRP14 (11525; 1:2,000 dilution) antibodies were obtained from Proteintech. The αFLAG (3165; 1:10,000 dilution), and αTubulin (T9026; 1:10,000 dilution) antibodies were obtained from Sigma.

### Reverse transcription – quantitative real-time PCR (RT-qPCR)

Total RNA was purified from transfected HeLa-HA cells using a RNeasy Micro RNA extraction Kit (Qiagen) with on column DNaseI treatment (Qiagen) and the RNA was eluted using 14 μL of DNA-free distilled deionized H_2_O. First strand cDNA synthesis was performed using a SuperScript III First Strand Synthesis kit (Invitrogen) using the supplied oligo dT primer according to the manufacturer’s instructions. The cDNA synthesis reactions were incubated at 50°C for 50 minutes and then heat inactivated at 80°C for 5 minutes and then diluted with H_2_O (1:1) before proceeding to qPCR.

Following cDNA synthesis, qPCR reactions (20 μL total volume) were carried out according to the manufacturer’s protocol, in triplicate for each condition, using the PowerUp SYBR Green Master Mix (Thermo Fisher Scientific) by combining 10 μL PowerUp SYBR Green Master Mix, forward (NeoTet1s: 5′-AGTCCCTTCCCGCTTCAGTGACAAC-3′) and reverse (NeoTet1as: 5′-CCTCGGCCTCTGAGCTATTC-3′) oligonucleotide primers (5 μM each), and 1 μL of diluted cDNA (see above). GAPDH oligonucleotide primers (forward: 5′-ACCACAGTCCATGCCATCAC-3′; reverse: 5′-TCCACCACCCTGTTGCTGTA-3′) were used as the internal control. RT-qPCR was carried out an ABI 7300 Real-Time PCR system (Thermo Fisher Scientific) using the following thermal cycling conditions: initial cycle of 50°C for 2 minutes; 95°C for 2 minutes; followed by 40 cycles of 15 seconds at 95°C, 15 seconds at 53°C, and 60 seconds at 72°C. The T method (46) was used to calculate relative *Alu* RNA expression levels using GAPDH RNA expression as the internal control. RNA expression data for each co-transfection condition is normalized to HeLa-HA co-transfected with TMO2F3 and *AlumneoI* and are the averages of three independent experiments.

## RESULTS

### Retrotransposition of an *Alu*Y element transcribed by an RNA pol II CMV promoter

To identify *Alu* sequences essential for retrotransposition, we employed a well-established cell culture-based *Alu* retrotransposition assay (4), where an *Alu*-permissive cell line (*i.e.*, HeLa-HA) (30,47) is co-transfected with a “driver” plasmid that expresses an epitope tagged version of L1 ORF2 (*i.e.*, TMO2F3) and a “reporter” plasmid (*e.g.*, *Aluneo*^Tet^) that expresses an *Alu* element that is tagged at its 3′ end with a retrotransposition indicator cassette (**Fig S1A**). The *Aluneo*^Tet^ plasmid expresses an *Alu*Y consensus RNA using an RNA pol III promoter that consists of the A and B boxes located within the *Alu*Y left monomer and a 66 bp sequence derived from the *7SL* RNA gene enhancer located upstream of the first *Alu*Y nucleotide (4,48,49). The retrotransposition indicator cassette (*neo*^Tet^) lacks a stretch of more than 3 thymidines and contains an antisense copy of the *neo* gene interrupted by a sense strand *Tetrahymena* group I self-splicing intron (4,33). An SV40 promoter sequence is located at the 5′ end of the *neo* gene, which drives *neo* expression after integration into genomic DNA. The *neo*^Tet^ indicator cassette is followed by a 44 bp poly(A) sequence followed immediately by an RNA pol III terminator sequence of 4 thymidines. Upon transcription, the group I intron undergoes self-splicing, reinstating the *neo* gene, and the resultant *Aluneo*^Tet^ transcript is inserted into genomic DNA by ORF2p, which confers G418 resistance to the cells.

The expression of *Alu* RNA from the *Aluneo*^Tet^ plasmid relies on an intact internal *Alu* pol III promoter. Therefore, mutations that disrupt the internal *Alu* promoter will negatively impact *Alu* transcription, making it difficult to determine how mutations or large deletions within *Alu* affect retrotransposition (25). To address this limitation, we designed *Alu* reporter vectors that utilize a constitutive RNA polymerase II (pol II) cytomegalovirus (CMV) immediate/early promoter to drive *Alu* RNA expression. We reasoned that a pol II promoter would facilitate the expression of extensively modified *Alu* sequences thereby allowing us to directly test the effects of large deletions within the *Alu* sequence on retrotransposition.

To construct plasmid vectors that express *Alu* RNA from a CMV promoter (*AlumneoI*) (**Fig S1A**), the *Alu*Y sequence from *Aluneo*^Tet^ was inserted into a pCEP4 mammalian expression vector downstream of the CMV promoter so that transcription would initiate at the first *Alu*Y nucleotide (**Fig S1B**). The CMV-driven *Alu*Y sequence was tagged with a pol II-compatible *mneoI* retrotransposition indicator cassette (34,50), which consists of an antisense copy of the neomycin phosphotransferase gene interrupted by a sense strand γ-globin intron sequence. An SV40 polyadenylation signal (PAS) sequence follows the *mneoI* indicator cassette. The retrotransposition of *AlumneoI* RNA confers G418-resistance to the HeLa-HA cells and allows retrotransposition activity to be quantified by counting the resultant G418-resistant foci (**Fig S1A**).

To determine if *Alu*Y could retrotranspose when expressed from a CMV promoter we co-transfected HeLa-HA cells with TMO2F3 and *AlumneoI* (**Figs 1B and S1A**). Retrotransposition assays revealed that, as expected, HeLa-HA cells that were co-transfected with TMO2F3 and *Aluneo*^Tet^ gave rise to high numbers of G418-resistant foci (∼390 foci on average per single well of a 6-well plate) (**Fig 1B**). Remarkably, co-transfection of HeLa-HA with TMO2F3 and *AlumneoI* resulted in approximately ∼90% of the number of G418-resistant foci compared to co-transfection with TMO2F3 and *Aluneo*^Tet^ (**Figs 1B** and **1C**). As a negative control, co-transfection with TMO2F3 and a version of *AlumneoI* that lacked the CMV promoter (Δ*AlumneoI*) did not give rise to G418-resistant foci (**Fig 1B**). Additional retrotransposition assays revealed that co-transfection with TMO2F3 and a version of *AlumneoI* that contained a blasticidin resistance retrotransposition indicator cassette (*Alublast*) (37) in place of the *mneoI* indicator also retrotransposed efficiently in HeLa-HA cells, demonstrating that retrotransposition of pol II-transcribed *Alu* RNA in HeLa-HA cells is not peculiar to the *mneoI* retrotransposition indicator cassette or to the selection conditions used in the assay (**Fig S1C**). As expected, additional control retrotransposition assays showed that HeLa-HA cells co-transfected with a version of TMO2F3 that contains a D702A missense mutation in the RT domain of ORF2 (TMO2F3^D702A^) and either *AlumneoI* or *Aluneo*^Tet^ only gave rise to single G418-resistant foci (**Fig S1D**). Control western blot experiments indicated that ORF2p-FLAG was expressed at similar levels in HeLa-HA cells co-transfected with TMO2F3 and each of the different *Alu*-expressing plasmids (**Fig 1D**).

We next used inverse PCR (see Methods) to determine whether the G418-resistant foci obtained from HeLa-HA cells co-transfected with TMO2F3 and *AlumneoI* contained structural hallmarks of L1-mediated retrotransposition events. *A bona fide* L1-mediated insertion is distinguished by three sequence features: flanking short variable length target site duplications (TSDs); a 3′ poly(A) tract; and integration into an L1 ORF2p endonuclease (L1 EN) consensus cleavage sequence resembling (5′-TTTT/A-3′, where the ′/′ indicates the cleavage site) (37,51–55). The analysis of two *AlumneoI* insertions (**Fig 1E**; c1983 and c2177) revealed the expected L1 TPRT signatures, as both *AlumneoI* insertions were: (**1**) flanked by short target site duplications of 15 and 11 bps, respectively; (**2**) ended in poly(A) tracts of ∼90-20 bps, respectively; and (3) integrated into an ORF2p L1 EN cleavage site (5’-TTTC/A-3’ and 5’-TTTT/A-3’), respectively. Notably, both insertions were full-length and contained an additional untemplated 5′ “G” nucleotide, which could have resulted from untemplated nucleotide addition by ORF2p terminal transferase activity during reverse transcription (35,56). Thus, these data indicate that the *AlumneoI* insertions resulted from L1 ORF2p-mediated retrotransposition.

### Embedded *Alu* sequences promote the retrotransposition of an RNA pol II transcript in HeLa-HA cells

Most cellular *Alu* RNAs are transcribed as part of larger pol II transcripts and are referred to as embedded *Alu* elements (1). Some studies suggest that embedded *Alu* elements facilitate SINE-VNTR-*Alu* (SVA) retrotransposition (57,58) and may contribute to processed pseudogene formation (59). To directly test whether embedded *Alu* elements promote the retrotransposition of pol II transcripts, we first co-transfected HeLa-HA cells with TMO2F3 and a reporter plasmid (EGFP*mneoI*) (30) that expresses the EGFP open reading frame tagged with an *mneoI* retrotransposition indicator. As expected, retrotransposition assays revealed that HeLa-HA cells co-transfected with TMO2F3 and EGFP*mneoI* did not give rise to G418-resistant foci, which is consistent with our previous analyses (30,40) **(Figs 2A** and **2B)**. In contrast, HeLa-HA that were co-transfected with TMO2F3 and a modified version of EGFP*mneoI* (EGFP*AlumneoI)* where an *Alu*Y element is inserted 826 bp downstream from the CMV transcription at the 3′ flank of the EGFP coding sequence gave rise to ∼10% the number of G418-resistant foci compared to cells transfected with TMO2F3 and *AlumneoI* **(Figs 2A** and **2B)**.

**Figure 2:**
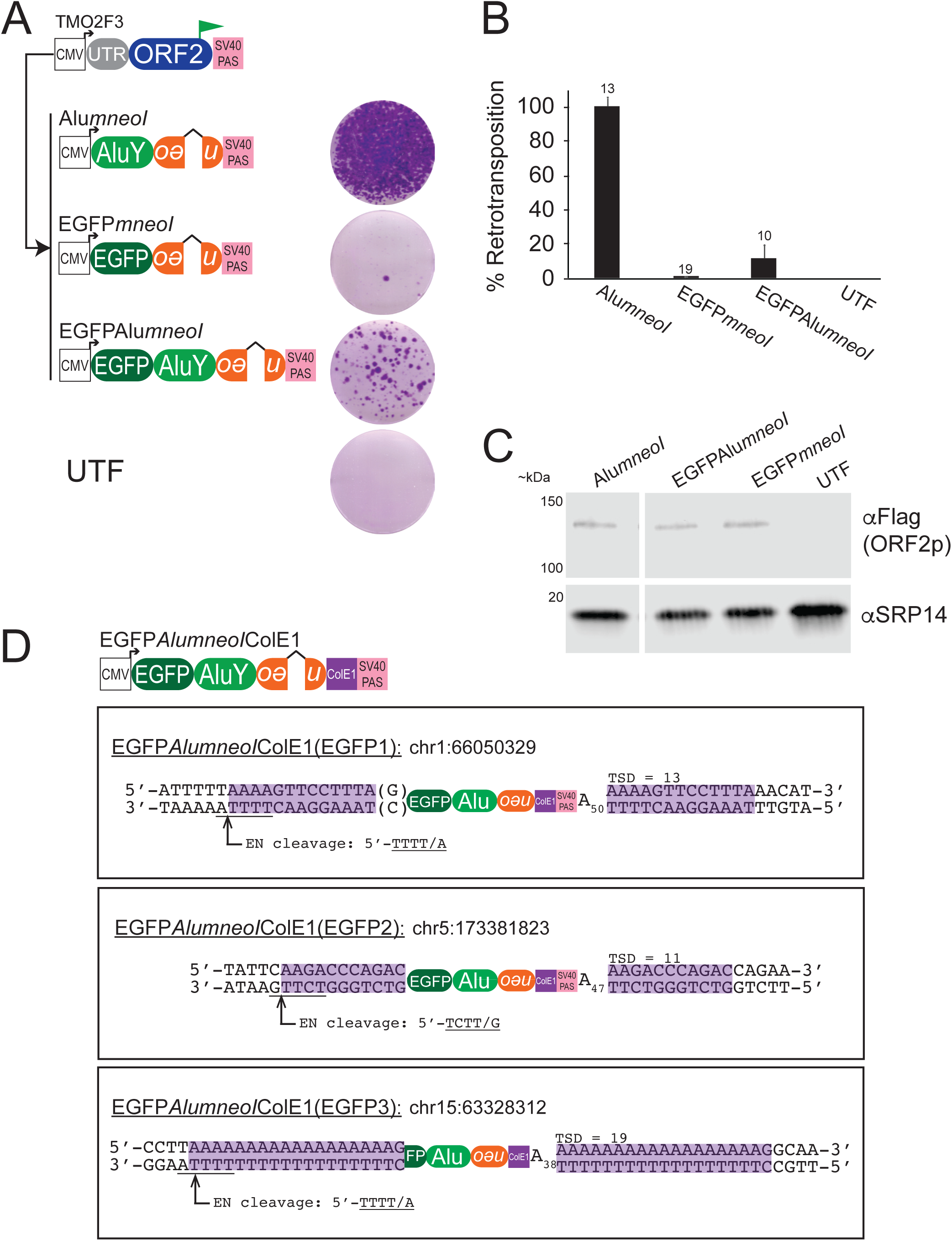
Embedded *Alu* sequences promote the retrotransposition of RNA pol II transcripts. *(A) Results of retrotransposition assays*. HeLa-HA cells were co-transfected with TMO2F3 and the indicated embedded *Alu* construct. Displayed to the right of construct are single wells of a representative six-well tissue culture plate from retrotransposition assays. *(B) Quantification of retrotransposition assays.* The X-axis indicates the *Alu* construct co-transfected with TMO2F3, and the Y-axis indicates average percent (%) retrotransposition normalized to *AlumneoI* for each transfection condition; error bars indicate standard deviation; (n) number of biological replicates indicated above error bars. *(C) ORF2p-Flag expression levels.* Western blots of HeLa-HA lysates co-transfected with TMO2F3 and the *Alu* retrotransposition construct (top of lane). The antibodies used for detection are indicated on the right and MW markers are indicated on the left of the blot image. Western blot experiments were repeated twice with similar results. *(D) Top panel: Diagram of the EGFPAluColE1 rescue vector.* Insertions were characterized from G418-resistant HeLa-HA cells that were co-transfected with TMO2F3 and EGFP*Alu*ColE1 using the rescue method (see Methods). *Bottom panel: Structure of three EGFPAluColE1 insertions (EGFP1, EGFP2, EGFP3).* Genomic insertion site locations are based on human reference genome sequence T2T CHM13v2.0/hs1. The EGFP*Alu*ColE1 insertions contain typical TPRT structural hallmarks indicative of L1 ORF2p-mediated retrotransposition (see main text). Target site duplications are highlighted in magenta. The L1 ORF2p EN target site is underlined; the arrow and “/” indicates the L1 EN cleavage site. Parentheses indicate an untemplated nucleotide.

To verify that EGFP*AlumneoI* insertions events were generated by ORF2p-mediated retrotransposition, we adapted a previously described method to directly characterize retrotransposition events and their associated flanking genomic DNA sequences as autonomously replicating, kanamycin-resistant plasmids in bacteria (see Methods) (35,44). Briefly, HeLa-HA cells were co-transfected with TMO2F3 and a EGFP*AlumneoI*ColE1 reporter plasmid, which is a modified version of EGFP*AlumneoI* that contains a ColE1 bacterial origin of replication located downstream of the *mneoI* indicator (35,44) (**Fig 2D**). Rescue of three EGFP*AlumneoI*ColE1 insertions (**Fig 2D**; EGFP1, EGFP2, and EGFP3) from G418-resistant foci revealed that each insertion was inserted into an ORF2p EN target site, contained short TSDs ranging from 13-19 bps, and had a poly(A) tract ranging from ∼38-50 nts in length (**Fig 2D**). Two insertions (EGFP1 and EGFP2) were full-length, and EGFP1 contained an untemplated “G” nucleotide at its 5′ end. EGFP3 was 5′-truncated within the *EGFP ORF* and truncated at its 3′ end prior to the SV40 PAS, which could be due to polyadenylation occurring at one of two cryptic PAS signals (AATAAA) located upstream of the SV40 PAS in the EGFP*AlumneoI*ColE1 plasmid. Thus, we conclude that EGFP*AlumneoI*ColE1 insertions resulted from L1-mediated retrotransposition and that embedded *Alu* sequences enhance the retrotransposition of RNA pol II transcripts in our assays.

### Identification of a minimal *Alu* domain RNA that associates with SRP9/14 in HeLa-HA cells

The ribosome association model of *Alu* retrotransposition hypothesizes that *Alu* RNA binds to SRP9/14 and localizes to ribosomes, which enables the encoded *Alu* RNA poly(A) tail to gain access to L1 ORF2p (4,19,25,26). Previous studies have demonstrated that point mutations within *Alu* that decrease binding to SRP9/14 also negatively impact *Alu* retrotransposition (25,29). Thus, the ability of *Alu* RNA to bind SRP9/14 is likely essential for *Alu* retrotransposition.

To test the above model, we first sought to identify the minimal *Alu* domain that associates with SRP9/14 in HeLa-HA cell extracts, reasoning that SRP9/14 binding is a prerequisite of *Alu* retrotransposition. Previous studies demonstrated that minimal *7SL*-derived *Alu* domain RNAs ranging between 47 and 86 nucleotides were sufficient to bind to SRP9/14 *in vitro* (60–62). We designed similar minimal *Alu* domain RNAs based on these studies (60–62) (**Fig 3A and Table S1**) and tested whether these minimal *Alu* domain RNAs could associate with SRP9/14 in HeLa-HA cells using RNA pull-down assays (**Fig 3B**). Briefly, 5′-biotin labeled *Alu* domain RNA oligonucleotides (**Fig 3B**) were synthesized, attached to magnetic streptavidin beads, and incubated in HeLa-HA cell lysates (see Methods). Proteins associated with the oligonucleotide then were analyzed by western blot using antibodies specific to SRP9 and SRP14.

**Figure 3:**
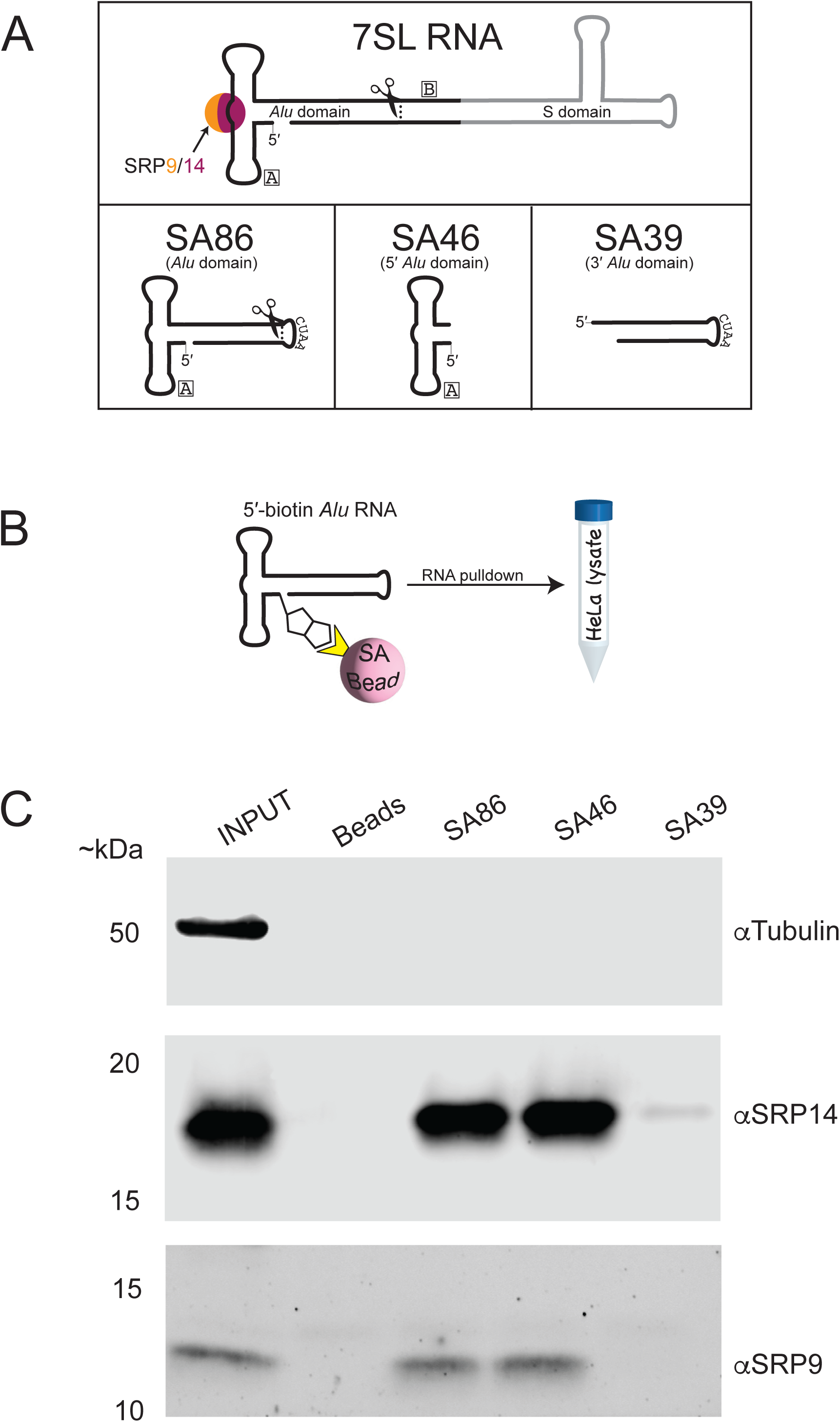
5′ *Alu* domain RNA binds to SRP9/14 in HeLa-HA cells. *(A) Diagram of Alu domain RNAs.* The *Alu* domain (black) of the *7SL* RNA (top) binds SRP9/14. SA86 (*7SL RNA Alu* domain) is an 86 nt fragment derived from the *7SL* RNA *Alu* domain where the scissors indicate the approximate location of SA86 in relation to the *7SL* RNA. The 3′ stem of SA86 is joined by a “CUAA” sequence. SA46 (5′ *Alu* domain) is composed of nts 1-46 of SA86. SA39 (3′ *Alu* domain) is composed of the last 39 nts of SA86. *(B) Schematic of Alu domain RNA pulldown experiments. Alu* domain RNAs containing a 5′ biotin moiety were pre-bound to streptavidin (SA) coated magnetic beads. *Alu* RNA-SA bead complexes were incubated in HeLa-HA cell lysates and western blotting was used to detect SRP9/14 eluted from *Alu* RNA-SA bead complexes. *(C) SRP9/14 binds to minimal Alu RNAs.* Western blots from RNA pulldown experiments. *Alu* RNA domain is indicated top of each lane. Input = ∼2% of WCL; Beads = empty beads (no RNA). The antibody used for detection is indicated on the right of the gel and MW markers indicated on the left of blot image. RNA pulldowns experiments were repeated three times with similar results.

RNA pull-down experiments performed in HeLa-HA lysates revealed that a synthetic *Alu* domain RNA oligonucleotide consisting of 86 nucleotides derived from the *7SL Alu* domain (SA86) associates with SRP9 and SRP14 in HeLa-HA cell extracts (**Fig 3C**). To further define the precise *Alu* domain sequences that are necessary to associate with SRP9/14 in HeLa-HA cells, we made further 3′ domain deletions of SA86 to generate a 46 nt 5′ *Alu* domain RNA (SA46) consisting of SA86 nucleotides 1-46 (**Fig 3A**). RNA pull-down experiments showed that SA86 and SA46 associate with similar levels of SRP9/14 in HeLa-HA cell extracts (**Fig 3C**). Pull-down experiments further revealed that the 3′ *Alu* domain (SA39), which consists of the 3′ domain of SA86 (nucleotides 47-86) (**Fig 3A**) failed to associate with SRP9/14 in HeLa-HA lysates (**Fig 3C**). An additional control demonstrated that SRP9/14 does not associate with empty streptavidin beads (**Fig 3C**). Thus, the minimal 5′ *Alu* domain RNA (SA46) is necessary and sufficient to associate with SRP9/14 in HeLa-HA cell extracts.

### Identification of a minimal *Alu* domain necessary and sufficient for retrotransposition

We next tested whether a minimal *Alu* domain is capable of retrotransposition by co-transfecting HeLa-HA cells with TMO2F3 and a reporter plasmid that uses a CMV promoter to drive expression of a minimal *Alu* domain tagged with an *mneoI* retrotransposition indicator cassette (**Fig 4A**). Retrotransposition assays revealed that HeLa-HA cells co-transfected with TMO2F3 and the 86 nt *7SL Alu* domain (SA86*mneoI*) gave rise to approximately 95% the number of G418-resistant foci compared to HeLa-HA cells co-transfected with TMO2F3 and *AlumneoI.* Inverse-PCR revealed that SA86*mneoI* insertions displayed canonical TPRT hallmarks (*i.e.*, the insertions were flanked by short variable length TSDs ranging in size between 10-18 bps, ended in a 3′ poly(A) sequences ranging between 59-82 A’s, and integrated into an L1 ORF2p endonuclease consensus cleavage sequence) (**Fig S2D**). The SA86*mneoI* (868B) insertion contained an untemplated 5′ “G” and the SA86*mneoI* (862A) insertion contained an untemplated 5′ “T”. Thus, an 86 nt *Alu* domain RNA (SA86) is mobilized by ORF2p-mediated retrotransposition in HeLa-HA cells.

**Figure 4:**
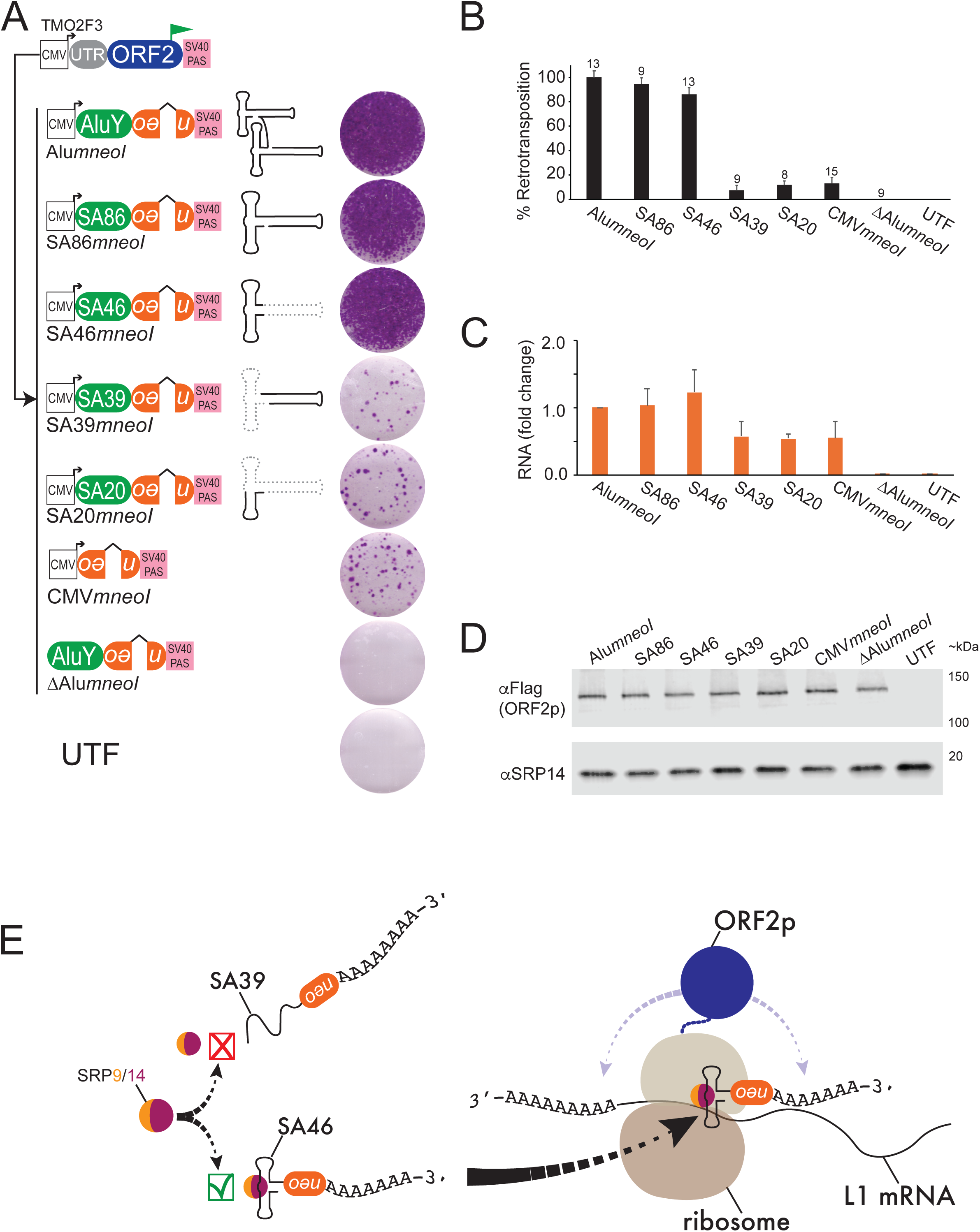
SA46 is necessary to promote efficient retrotransposition. *(A) Results of retrotransposition assays*. HeLa-HA cells were co-transfected with TMO2F3 and the indicated *Alu*-containing pol II transcribed RNA construct with a schematic diagram of the respective *Alu* domain RNA. Displayed next to each plasmid are single wells of a representative six-well tissue culture plate from retrotransposition assays. *(B) Quantification of retrotransposition assays.* The X-axis indicates the *Alu* RNA expressing construct co-transfected with TMO2F3, and the Y-axis indicates average percent (%) retrotransposition normalized to *Alu*mneoI for each transfection condition; error bars indicate standard deviations; (n) number of biological replicates are indicated above the error bars. *(C) Quantification of Alu domain RNA by RT-qPCR.* The X-axis indicates the *Alu* RNA expressing construct co-transfected with TMO2F3. The Y-axis indicates the RNA fold change normalized to *AlumneoI* from three independent experiments for each transfection condition with the exception of ΔCMV*Alu*mneoI, which is based on two independent experiments; error bars indicate the standard deviations. *(D) ORF2p-Flag expression levels.* Western blots of HeLa-HA lysates co-transfected with TMO2F3 and the *Alu* retrotransposition construct (indicated top of each lane). Western blot experiments were done twice with similar results. The antibody used for detection is indicated on right of the blot image and MW markers are indicated on the left of blot image. *(E) Model of SA46 retrotransposition. Alu* RNAs that contain an intact 5′ *Alu* domain (*i.e.*, *Alu*Y, SA86, SA46) bind SRP9/14 to form a stable *Alu* RNP that can localize to a ribosome where the *Alu* encoded poly(A) tract can compete with the L1 poly(A) tail to bind ORF2p. Polyadenylated RNA transcripts that do not contain a stable *Alu* domain (*i.e.*, SA39, SA20, CMV*mneoI*) cannot efficiently bind SRP9/14, may be less stable compared to 5′ *Alu* domain containing RNAs, and likely do not associate with ribosomes. Thus, SA39 and SA20 have a much lower propensity for retrotransposition when compared to *Alu*Y, SA86 and SA46.

To define the minimum *Alu* domain sequence required for retrotransposition, HeLa-HA cells were co-transfected with TMO2F3 and the 46 nt 5′ *Alu* domain (SA46*mneoI*), which gave rise to approximately 86% the number of G418-resistant foci compared to HeLa-HA cells co-transfected with TMO2F3 and *AlumneoI* (**Figs 4A** and **4B**). Similarly, HeLa-HA cells co-transfected with TMO2F3 and a construct expressing nucleotides 2-47 of the consensus *Alu*Y element left monomer 5′ *Alu* domain (AY46 *mneoI*) gave rise to approximately 73% the number of G418-resistant foci compared to HeLa-HA cells co-transfected with TMO2F3 and *AlumneoI* (**Figs S2A and S2B**). In contrast, transfection of HeLa-HA cells with TMO2F3 and the 3′ *Alu* domain (SA39*mneoI)* gave rise to only 7% the number of G418-resistant foci when compared to HeLa-HA cells co-transfected with TMO2F3 and *AlumneoI* (**Figs 4A** and **4B**). Similarly, HeLa-HA cells co-transfected with TMO2F3 and a 5′ *Alu* domain deletion mutant that consists only of the first 20 nucleotides of SA86 (SA20*mneoI*) gave rise to 12% the number of G418-resistant foci when compared to HeLa-HA cells co-transfected with TMO2F3 and *AlumneoI* (**Figs 4A** and **4B**). As expected, control retrotransposition assays revealed that HeLa-HA co-transfected with TMO2F3 and Δ*AlumneoI* did not give rise to G418 resistant colonies (**Figs 4A** and **4B**). Thus, a 46 nt 5′ *Alu* domain is required to mediate efficient retrotransposition in HeLa-HA cells under the conditions of our assay.

To determine whether L1 ORF2p preferentially mediates insertion of the *Alu* domain RNAs compared to non-*Alu*-containing RNA transcripts, we co-transfected HeLa-HA cells with TMO2F3 and a plasmid that uses CMV to express the *mneoI* indicator cassette and lacks any *Alu* RNA sequences (CMV*mneoI*). Retrotransposition assays revealed that HeLa-HA cells co-transfected with TMO2F3 and CMV*mneoI* gave rise to 13% the number of G418-resistant foci compared to co-transfection with TMO2F3 and *AlumneoI* (**Figs 4A** and **4B**, see Discussion). Similarly, HeLa-HA cells co-transfected with TMO2F3 and a plasmid that uses the CMV promoter to express a blasticidin-resistance (*mblastI*) retrotransposition indicator cassette that lacks *Alu* RNA sequences gave rise to comparable levels of blasticidin resistant foci (data not shown).

To determine whether CMV*mneoI* insertions resulted from ORF2p-mediated retrotransposition, we co-transfected HeLa-HA with TMO2F3 and a version of CMV*mneoI* that also contains a contains a ColE1 bacterial origin of replication located after the *mneoI* indicator (35,44). We then characterized CMV*mneoI*ColE1 insertion events from G418-resistant cells, which confirmed their identity as L1-mediated retrotransposition events (*i.e.*, the insertions [581 and 386] were flanked by 15 and 16 bp TSDs, respectively, ended in ∼47 and 60 bp 3′ poly(A) sequences, respectively, and integrated into an L1 ORF2p endonuclease consensus cleavage sequence) (**Fig S2E**). Thus, the data suggest L1 ORF2p preferentially retrotransposes RNAs containing the 5′ *Alu* domain compared to other polyadenylated transcripts. Intriguingly, these results may further suggest that short non-coding polyadenylated transcripts may be preferentially retrotransposed when compared to polyadenylated transcripts that contain an intact open reading frame (compare **Fig 4A to Fig 2A**; EGFP*mneoI* results and see Discussion section).

To test whether the differences in the retrotransposition of the *Alu* domain RNA reporter constructs resulted from differences in the steady state reporter RNA levels, we performed real-time quantitative PCR (RT-qPCR) using primers specific to the *mneoI* reporter cassette. These experiments demonstrated that, in HeLa-HA cells co-transfected with TMO2F3 and either *AlumneoI,* SA86*mneoI*, or SA46*mneoI*, *mneoI* expression levels were similar after approximately 48 hours post-transfection (**Fig 4C**). By comparison, *mneoI* expression levels were approximately 50% less in HeLa-HA cells co-transfected with TMO2F3 and SA39*mneoI,* SA20*mneoI*, or CMV*mneoI*. These lower RNA levels could reflect slight decreases in the stability of transcripts that lack the *Alu* domain. That being stated, the lower RNA levels cannot explain the ∼80-90% reduction in retrotransposition. Control western blot experiments further confirmed that ORF2p was expressed at similar levels in HeLa-HA cells co-transfected with TMO2F3 and each of the above reporter plasmids ∼48 hours post-transfection (**Fig 4D**). Thus, our data indicate that the *Alu* 5′ domain is required to mediate a single round of retrotransposition in HeLa-HA cells.

## DISCUSSION

Here, we identified a minimal *Alu* domain RNA consisting of 46 nts that is necessary to facilitate a single round of *Alu* retrotransposition in HeLa-HA cells. The fact that *Alu* transcription by pol III requires an intact internal A and B box has previously made it difficult to precisely define what *Alu* sequences, besides the poly(A) tail, are necessary for retrotransposition using previous *Alu* retrotransposition assays (4) because deletion or mutation of sequences encompassing the A and/or B box would negatively impact *Alu* transcription, and hence retrotransposition (25). To overcome this limitation, we constructed plasmid vectors that express *Alu* elements from a CMV pol II promoter and used an SV40 PAS for the post-transcriptional addition of a 3′ poly(A) tract to the *Alu* RNA. Our data indicate that *Alu* RNA expressed from a pol II promoter faithfully recapitulates *Alu* retrotransposition (**Figs 1B, 2A, and 3A**), and pol II-transcribed *Alu* insertions exhibit the hallmarks of L1 ORF2p-mediated retrotransposition events (**Figs 1E, S2B, S2D, S2D, and S2E**). Notably, we also constructed reporter vectors that use an external U6 promoter to drive *Alu* domain RNA expression, which resulted in efficient retrotransposition when co-transfected with TMO2F3 (data not shown). Thus, other external promoters besides CMV can be used to drive SINE RNA expression that result in retrotransposition. We envision that our CMV pol II *Alu* expressing vectors could be adapted to express other SINE RNAs (*e.g.*, *tRNA*-derived and *5S ribosomal RNA* SINEs) (63–66) and could easily be modified with other available retrotransposition indicators (**Fig S1C**) (38,67,68), to identify domains required for retrotransposition of other pol III transcribed SINE RNAs.

Our results highlight the importance of *Alu* RNA structure in mediating efficient *Alu* retrotransposition. We demonstrated that the 5′ *Alu* domain (SA46 and AY46) is required to mediate efficient *Alu* retrotransposition in HeLa-HA cells. It is likely that the 46 nt 5’ *Alu* RNA domain folds into a structure that facilitates SRP9/14 binding and interaction with the ribosome (**Fig 4E**) (4,19,25). RNAs that lacks a properly folded *Alu* domain, such as SA39 or SA20, likely exhibit decreased retrotransposition because they fail to efficiently bind SRP9/14, which hamper their ability to localize to ribosomes and compete for ORF2p binding (**Fig 4E**). We speculate that tRNA-derived SINEs, and perhaps other SINE RNAs, have adapted similar strategies to gain access to ORF2p (30). How these different non-coding SINE RNAs retrotranspose and whether they compete with *Alu* for host factors and/or ribosome binding requires elucidation. However, it is likely that RNA structure plays a significant role in the retrotransposition for all SINEs (25,69).

The SA46 and SA86 *Alu* domains are derived from the *7SL* RNA (60,61). The 5′ *7SL Alu* domain (SA46) and the consensus *Alu*Y 5′ left monomer *Alu* domain (AY46) differ by seven nucleotides and share ∼85% sequence identity (**Fig S2C**). Our data showed that despite these differences, AY46 exhibited high levels of retrotransposition that are comparable to SA46 (**Figs S2A and S2B**). The nucleotide differences between SA46 and AY46 occur predominantly in unpaired loop regions within the 5′ *Alu* domain secondary structure (**Fig S2C**), which tend to be less conserved amongst eukaryotic SRP RNAs (25). Notably, because SA46 and other truncated 5′ *Alu* RNAs do not contain intact RNA pol III promoters they are incapable of additional rounds of retrotransposition unless they fortuitously insert into a genomic context that provides both a promoter and transcription termination signal. This scenario is reminiscent of previously reported Spliced Integrated Retrotransposable Elements (SpIRE), U6/L1 chimeric pseudogenes, and processed pseudogene retrotransposition events, which can undergo an initial round of retrotransposition, but then are rendered inactive due to a loss of promoter sequences (40,45,70–74). Although the 3′ *Alu* domain (SA39) is not needed for efficient retrotransposition, it is necessary in the context of an active *Alu* retrotransposon because it harbors *Alu* promoter sequences (*i.e.*, the B box), which are necessary for *Alu* transcription. It will be interesting to determine whether the *Alu* 3′ domain sequences have any additional functions in *Alu* retrotransposition.

Our data demonstrate that embedded *Alu* sequences could facilitate the retrotransposition of pol II RNA transcripts (**Figs 2A** and **2D**). Our functional data is consistent with a previous bioinformatics study, which suggested that 3′ UTR-embedded *Alu* elements might facilitate the formation of processed pseudogenes (59). Consistently, the presence of a 3′-transduced *Alu* element is reported to promote SVA retrotransposition and could have contributed to the rapid expansion of the SVA_F1 subfamily (57,75). Embedded *Alu* sequences also have been shown to play a role in processes such as alternative splicing, RNA transcript localization, and RNA editing by ADAR (adenosine deaminases acting on RNA) proteins (reviewed in: (1,76,77)). Lastly, previous studies have shown that SRP9/14 binds to larger RNA transcripts that contain embedded *Alu* sequences (22). Thus, we speculate that embedded *Alu* sequences offer the potential to fold into a structure that interacts with SRP9/14 to mediate ribosomal localization and retrotransposition of larger RNA transcripts that contain them.

The observation of relatively high levels of retrotransposition from CMV*mneoI*, SA20*mneoI*, and SA39*mneoI* reporters compared to the background rate of G418-resitant colony formation in untransfected HeLa-HA or HeLa-HA co-transfected with TMO2F3 and Δ*AlumneoI* (**Fig 4**) is notable considering the low rate of processed pseudogene formation reported in previous studies (4,40). Our reporter constructs (*i.e.*, CMV*mneoI*, SA20*mneoI*, SA39*mneoI*) are predicted to express non-coding RNA polyadenylated transcripts that do not encode long (>45 amino acids) open reading frames. By comparison, the retrotransposition rate of the EGFP*mneoI* reporter, which encodes the EGFP open reading frame, is closer to the estimated rate of processed pseudogene formation reported previously (40) (**see Fig 2**, ∼0.4% compared to *AlumneoI*). Thus, these data suggest that non-coding RNA transcripts could be preferred retrotransposition templates when compared to RNA transcripts that contain open reading frames (*i.e.*, processed pseudogene precursor RNAs), which is consistent with the fact that many SINEs are derived from non-coding RNAs (65,78,79).

In conclusion, we have identified a 46 nt 5′ *Alu* domain that binds to SRP9/14 and is required for highly efficient retrotransposition. Our findings are consistent with the ribosomal localization model of *Alu* retrotransposition (4,19,25,26), which explains how *Alu* RNA is preferentially retrotransposed compared to the myriad of other polyadenylated RNAs in the cell. After *Alu* is transcribed by pol III, the 46 nt 5′ *Alu* domain folds into a structure that facilitates binding to SRP9/14 to form an *Alu* RNP that interacts with the ribosome. The localization of *Alu* RNA to ribosomes places the *Alu* RNA poly(A) tail in the right place and time to compete with the L1 RNA poly(A) to co-translationally bind L1 ORF2p. It is remarkable that a 46 nt sequence derived from the ancient *7SL Alu* domain is responsible for helping to generate nearly 11% of the human genome (7). Indeed, it would be interesting to examine whether molecular mimics of this 46 nt 5′ *Alu* domain sequence could promote the enhanced retrotransposition of other cellular RNAs.

## ACKNOWLEDGEMENTS

We thank Dr. Thierry Heidmann for providing the p*Aluneo*^Tet^, and Dr. Astrid Engel for providing the HeLa-HA strain. We also thank Dr. Mitsuhiro Nakamura, Dr. Jacob Mueller, and J.V.M. lab members for critical reading of this manuscript and helpful discussions during the course of this study. We are grateful to Henrietta Lacks, now deceased, and to her surviving family members for their contributions to biomedical research. The HeLa-HA cell line that was established from her tumor cells without her knowledge or consent in 1951 has made significant contributions to scientific progress and advances in human health.

## AUTHOR CONTRIBUTIONS

JBM and JVM designed the study. JBM and JY conducted experiments. JBM, JY, and JVM analyzed the data. JBM, JY, and JVM wrote and edited the manuscript.

## FIGURE LEGENDS

**Supplementary Figure S1.**
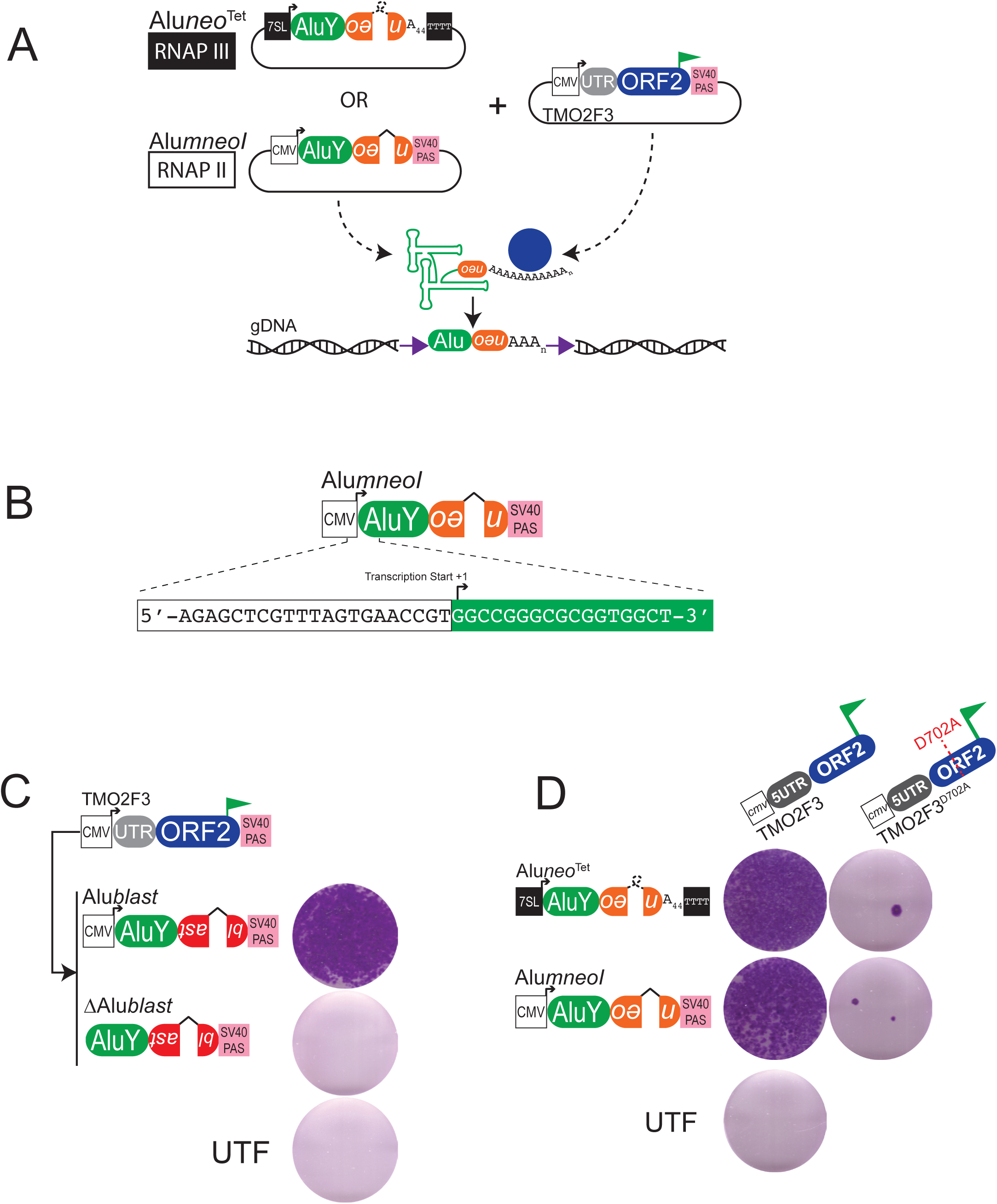
Related to Figure 1: *(A) Schematic of the Alu retrotransposition assay.* HeLa-HA cells are co-transfected with TMO2F3 and *Aluneo*^Tet^ or *AlumneoI.* TMO2F3 expresses ORF2p with a carboxyl terminus 3XFLAG tag (green flag). ORF2p expression is augmented by a CMV promoter (white square), the native L1 5′ UTR (grey oval), and an SV40 polyadenylation signal sequence (pink square). The *Aluneo*^Tet^ retrotransposition marker (orange oval) contains a backwards copy of the neomycin phosphotransferase gene interrupted by a self-splicing group I intron (loop) that is in the same transcriptional orientation as the *Alu* sequence. A 44 bp encoded poly(A) tract follows the *neo*^Tet^ sequence. *Aluneo*^Tet^ expression is augmented by a *7SL* gene enhancer (black square) and a sequence of four consecutive thymidine residues (black square) located downstream of the 44 bp poly(A) sequence. The *AlumneoI* retrotransposition marker (orange oval) contains a backwards copy of the neomycin phosphotransferase gene interrupted by a γ-globin intron (^) that is in the same transcriptional orientation as the *Alu* sequence. *AlumneoI* expression is augmented by a CMV promoter (white square) and an SV40 polyadenylation signal sequence (pink square) located after the *mneoI* indicator cassette. The neomycin phosphotransferase gene can only be expressed when the *Alu* transcript is spliced and subsequently is inserted into genomic DNA by L1 ORF2p (Blue circle). The resultant insertions are flanked by target site duplications (purple arrows). *(B) AlumneoI transcription start site.* Sequence of the *AlumneoI* plasmid at the junction between the CMV promoter (white box) and the *Alu*Y (green box, white letters) transcription start site. *(C) Retrotransposition results for Alublast.* Diagram of plasmids used in the assay. *Alublast* or an *Alublast* lacking the CMV promoter (Δ*Alublast)* were co-transfected with TMO2F3. To the right of the plasmids are qualitative retrotransposition results. Displayed are representative images from a 6-well plate. Experiments conducted at least three times with similar results. *(D) Retrotransposition results for Alu plasmids co-transfected with L1 ORF2p reverse transcriptase mutant.* Diagram of plasmids used in the assay. *Alu*neo^Tet^ or *AlumneoI* were co-transfected with an ORF2p “driver” plasmid that expresses wild type ORF2p (TMO2F3) or a reverse transcriptase mutant version of ORF2p (TMO2F3^D702A^). To the right of *Alu* plasmids are the qualitative retrotransposition results. Displayed are representative images from a 6-well plate of at least two independent experiments.

**Supplementary Figure S2.**
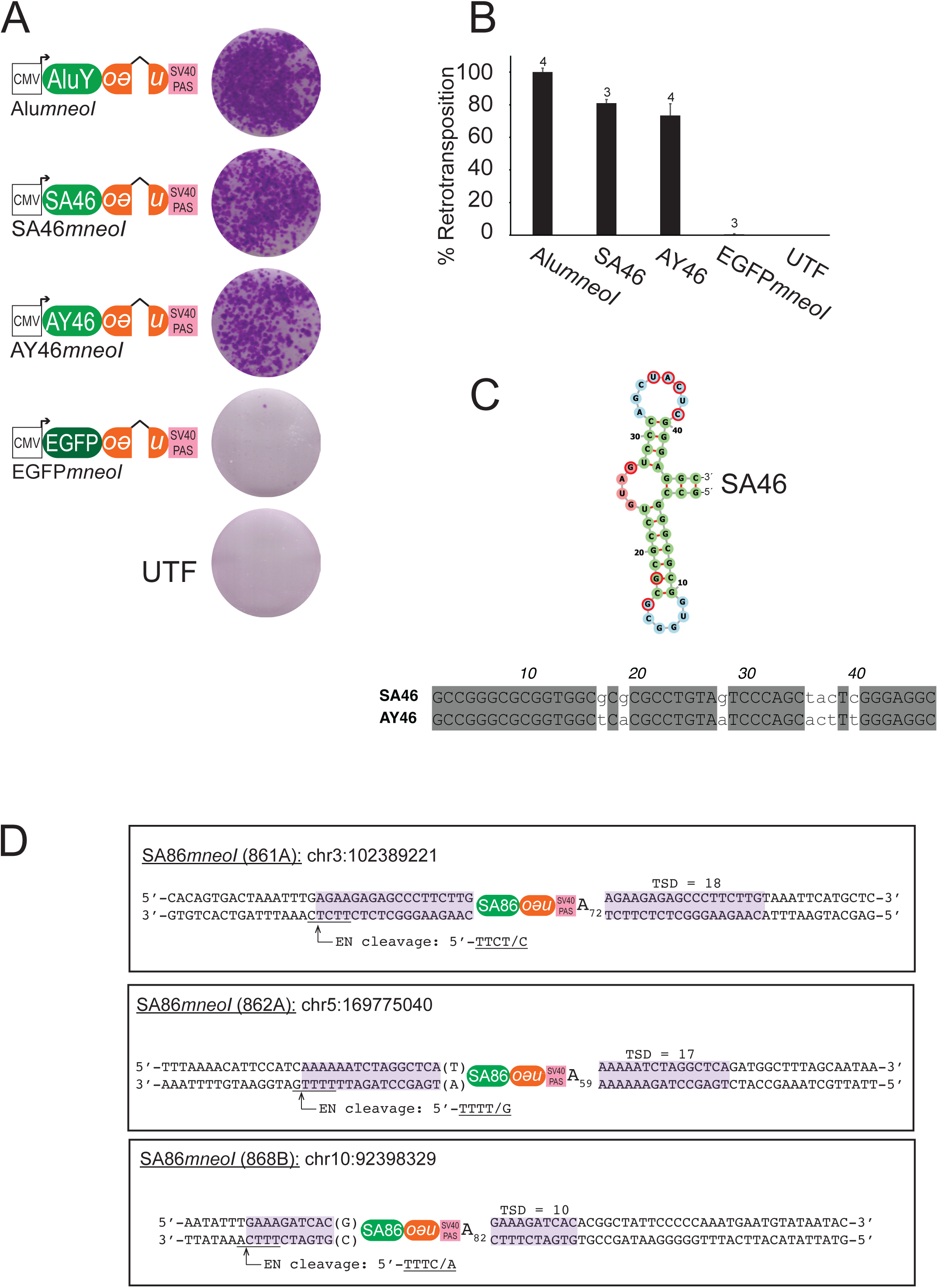
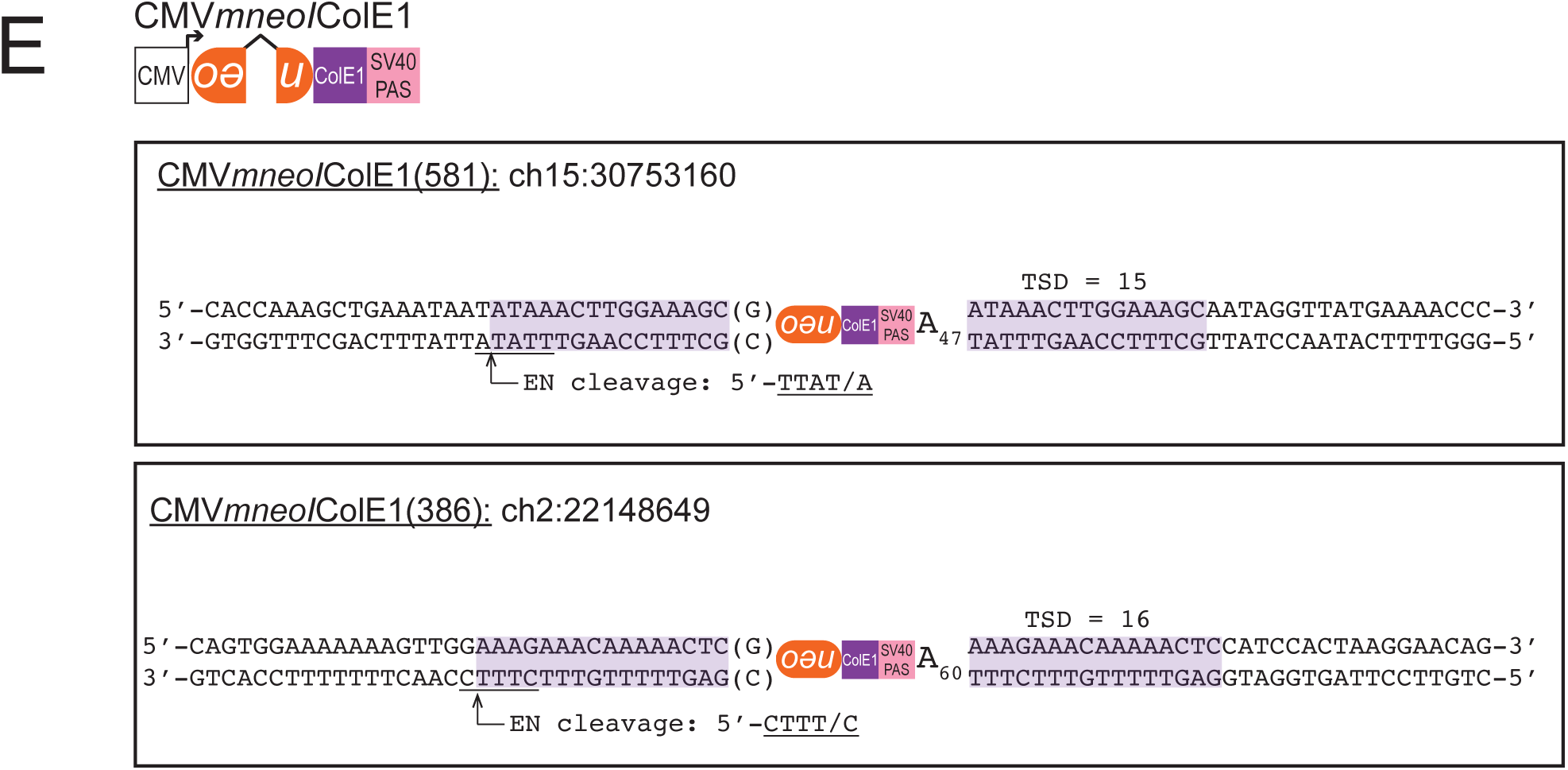
Related to Figure 4: *(A) AluY46 retrotransposition results*. HeLa-HA cells were co-transfected with TMO2F3 and the indicated *Alu* RNA construct. To the right of construct are displayed are single wells of a representative six-well tissue culture plate from retrotransposition assays. *(B) Quantification of retrotransposition assays.* The X-axis indicates the *Alu* RNA expressing construct co-transfected with TMO2F3. The Y-axis indicates the average percent (%) retrotransposition normalized to *AlumneoI* from (n) independent biological replicates for each transfection condition; error bars indicate standard deviations; (n) number of biological replicates indicated above error bars. *(C) Sequence comparison between SA46 and AY46.* (Top) Secondary structure of the SA46 5′ *Alu* domain (61). Nucleotide differences between SA46 and AY46 are outlined in red. (Bottom) Sequence alignment between SA46 and AY46. *(D) Structure of SA86mneoI insertions characterized by inverse PCR.* Genomic insertion site locations are based on human reference genome sequence T2T CHM13v2.0/hs1. The SA86*mneoI* insertions contain typical TPRT structural hallmarks indicative of L1 ORF2p-mediated retrotransposition (see main text). Target site duplications are highlighted in magenta. The L1 ORF2p EN target site is underlined; the arrow and “/” indicates the L1 EN cleavage site. Parentheses indicate an untemplated nucleotide present at the 5′ end of the insertion. *(E) Top panel: Diagram of CMVmneoIColE1 rescue vector.* Insertions were characterized from G418-resistant HeLa-HA cells that were co-transfected with TMO2F3 and CMV*mneoI*ColE1 using the rescue method (see Methods). *Bottom panel: Structure of de novo CMVmneoIColE1 insertions (581 and 386).* Insertions were characterized from G418-resistant HeLa-HA cells that were co-transfected with TMO2F3 and CMV*mneoI*ColE1 using the rescue method. Genomic insertion site locations are based on human reference genome sequence T2T CHM13v2.0/hs1. The insertions contain typical TPRT structural hallmarks indicative of L1 ORF2p-mediated retrotransposition (see main text). Target site duplications are highlighted in magenta. The L1 ORF2p EN target site is underlined; the arrow and “/” indicates the L1 EN cleavage site. Parentheses indicate untemplated nucleotides present at the 5′ end of the insertion.

**Supplementary Table S1.**
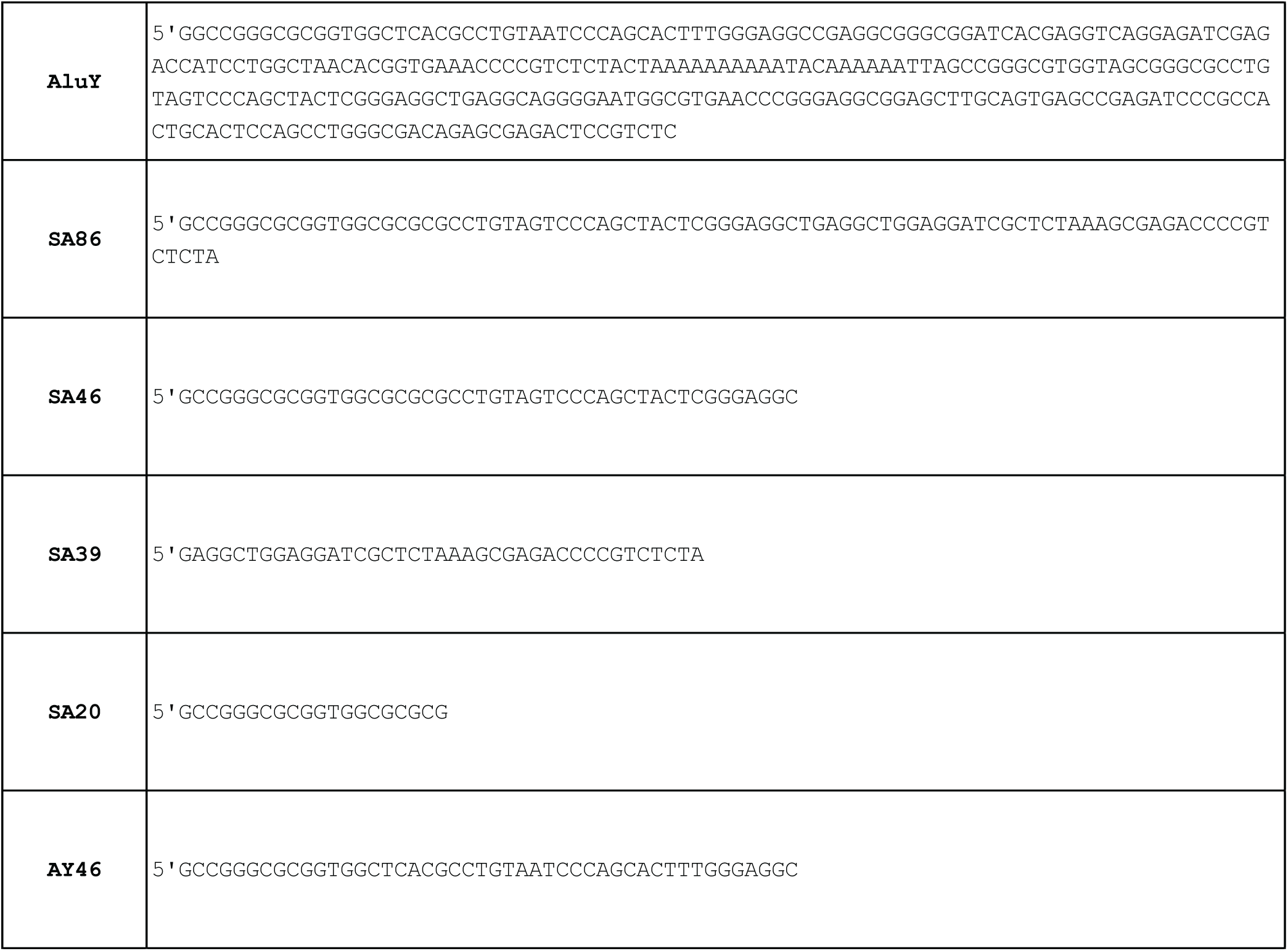
A list of *Alu* sequences used in this study.

